# Direct measurement of the mutation rate and its evolutionary consequences in a critically endangered mollusk

**DOI:** 10.1101/2024.09.16.613283

**Authors:** T. Brock Wooldridge, Sarah Ford, Holland Conwell, John Hyde, Kelley Harris, Beth Shapiro

## Abstract

The rate at which mutations arise is a fundamental parameter of biology. Despite recent progress in measuring germline mutation rates across diverse taxa, such estimates are missing for much of Earth’s biodiversity. We present the first estimate of a germline mutation rate from the phylum Mollusca, which is diverged by more than 1200 Ma years from the closest relative for which a mutation rate estimate exists. We sequenced three pedigreed families of the white abalone *Haliotis sorenseni*, a long-lived, large-bodied, and critically endangered mollusk, and estimated a *de novo* mutation rate of 8.60e-09 single nucleotide mutations per site per generation. This mutation rate is similar to rates measured in vertebrates with similar generation times and longevity to abalone, and higher than mutation rates measured in faster-reproducing invertebrates. We use our estimated rate to infer baseline effective population sizes (N_e_) across multiple Pacific abalone and find that abalone persisted over most of their evolutionary history as large and stable populations, in contrast to extreme fluctuations over recent history and small census sizes in the present day. We then use our mutation rate to infer the timing and pattern of evolution of the abalone genus *Haliotis*, which was previously unknown due to few fossil calibrations. Our results are an important step toward understanding mutation rate evolution and establish a key parameter for conservation and evolutionary genomics research in mollusks.

## Introduction

Mutations are the ultimate source from which variation arises. Although mutation is a fundamental feature of all life on Earth, the rate at which new mutations arise can vary considerably between and within species. Consequently, the mutation rate and the extent to which it is fine-tuned by natural selection has long held the interest of biologists (Sturtevant 1937; Lynch et al. 2016).

Mutations may occur in any cell type but only mutations that occur in an organism’s germline contribute to subsequent generations and drive evolutionary innovation (Bergeron et al. 2023). Within multicellular eukaryotes, germline mutation rates (GMRs) vary by at least three orders of magnitude, and the etiology of this variation is not completely understood (Lynch 2010). Mutation rates per generation are generally highest in large-bodied, long-lived organisms with modest effective population sizes, and several mechanisms (which are not mutually exclusive) have been proposed to explain this trend. Long generation times could increase GMRs by allowing more time for mutations to accumulate in spermatocytes and oocytes prior to reproduction. Consequently, the length of time elapsed between puberty and reproduction has been proposed to explain observed mutation rate variation among vertebrates (Thomas 2018). However, small effective population sizes could also explain this variation, since weakly deleterious alleles may drift to high frequencies in small populations, increasing observed mutation rates (Lynch 2010). Both generation time and effective population size are similarly correlated with mutation rate variation in empirical datasets, making it difficult to disentangle the relative etiological contributions of reproductive longevity and the drift-barrier effect (Wang and Obbard 2023).

Current understanding of the causes and extent of GMR variation is shaped by available direct estimates of germline mutation rates. However, these data are not representative of Earth’s biodiversity. Roughly 83% of animals with an estimated GMR are vertebrates (Wang and Obbard 2023), although that vertebrates represent only 4.6% of animal diversity (Bánki et al. 2024). Some animal phyla are entirely unrepresented among available data for GMRs, including Mollusca. Mollusks encompass a broad diversity of form and function, spanning terrestrial species like the common garden snail to the deep ocean dwelling giant squid, as well as considerable variation among lineages in population size, gamete production, parental investment, and longevity (Ponder and Lindberg 2008), and may therefore provide useful data to better understand the evolution of variability among GMRs. The closest relative of Mollusca with an estimated GMR, however, shares a common ancestor with mollusks over 600 million years ago, a distance that represents more than 1200 Ma of independent evolution (Fig. 1).

**Figure 1.**
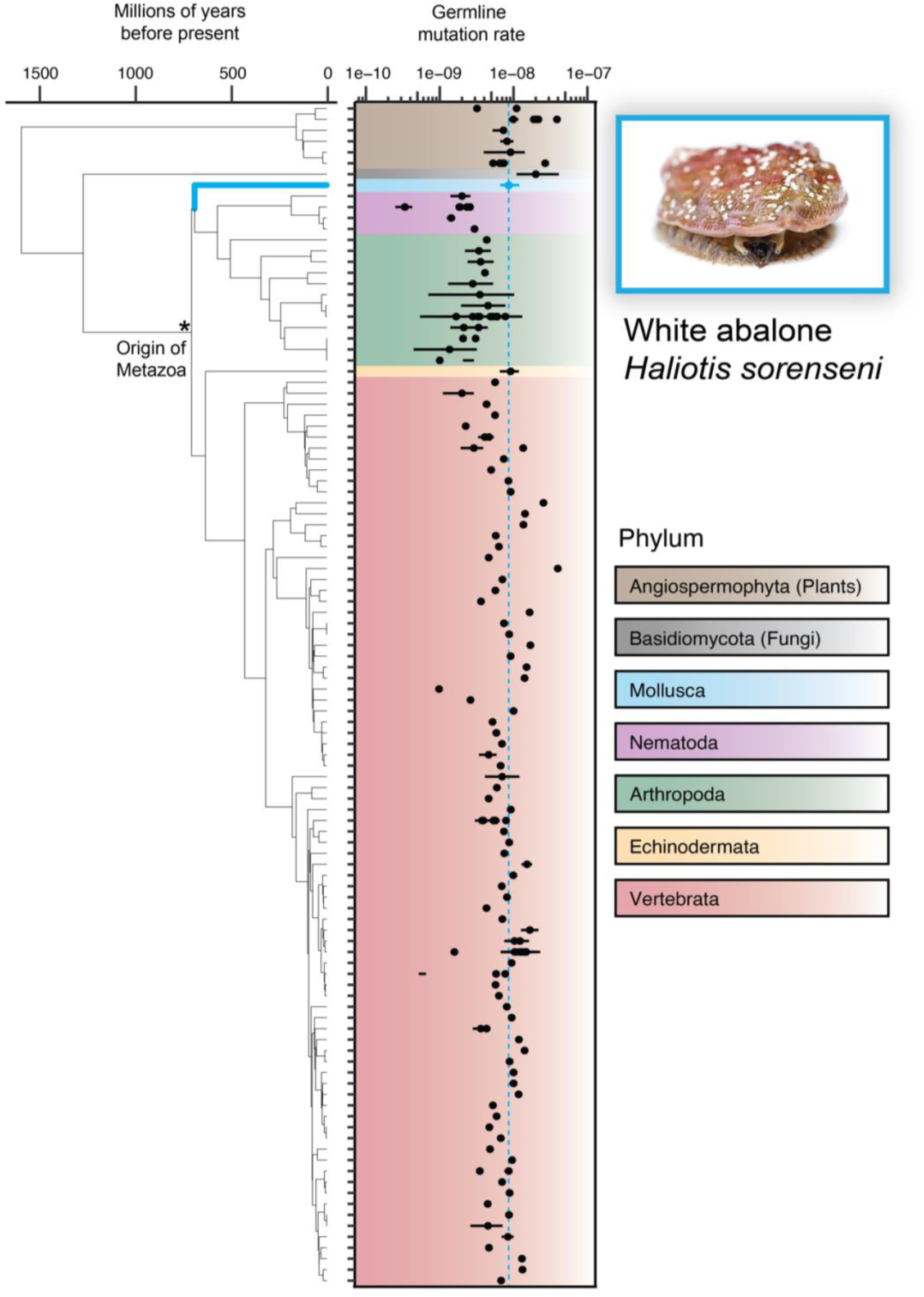
Distribution of all SNP germline mutation rate estimates for multicellular eukaryotes, adapted from Wang and Obbard et al. (2023). Time-scaled phylogeny to the left retrieved from *TimeTree*. White abalone (*Haliotis sorenseni*), the sole mollusk and focal species of this study, is highlighted in blue.

The rate at which mollusks accumulate *de novo* mutations remains unknown, although the high genetic diversity in many mollusk populations has led to speculation of a fast GRM (Zhang et al. 2012; Hoeh et al. 1996; Cutter, Jovelin, and Dey 2013; Launey and Hedgecock 2001). Previous work, for example, estimated a GMR for the Pacific oyster *Crassostrea gigas* that was 90 times faster than that of *Drosophila,* based on the oyster’s anomalously high deleterious mutation load (Plough, Shin, and Hedgecock 2016). This estimate was predicated on the theoretical relationship between the frequency of lethal mutations and mutation rates (Nei 1968), however, and not a direct estimate of the GMR (Plough, Shin, and Hedgecock 2016). Although fast mutation rates can lead to high levels of genetic diversity, genetic diversity (*π*) can also be maintained in populations with large effective sizes (N_e_) without requiring faster rates of mutation (*μ*) (*π* = 4N_e_μ (Nei and Tajima 1981). Many marine mollusks do have large census population sizes (N_c_) but effective size may be much smaller due to processes like ‘sweepstakes reproduction’, in which a handful of highly fecund individuals reproduce each generation (Hedgecock and Pudovkin 2011; Hedrick 2005). Therefore, uncertainty surrounding N_e_ makes inference of a GMR based on population genetic diversity difficult (Harrang et al. 2013).

In contrast to inferences based on population genetic diversity, phylogenetic analyses of mollusks have suggested lower GMRs. Based on rates of nucleotide substitution between mollusk lineages calibrated by fossil ages, these estimates are on the order of 3 x 10 ^-9^ mutations/bp/generation (A. Li et al. 2021; Allio et al. 2017).Phylogenetic approaches to estimating mutation rates are, however, error-prone due to uncertainty in fossil calibration, generation time, mutation saturation over long timescales, and difficulty in identifying truly neutral sites for estimation (Wang and Obbard 2023; Scally and Durbin 2012). Because indirect estimates of mutation rates in mollusks vary to such an extent, a direct estimate based on pedigree sampling is needed.

Accurate estimates of germline mutation rates have practical applications for conservation and evolutionary genomics research. Effective population size, which describes the size of an idealized Wright-Fisher population that would exhibit the magnitude of genetic drift and inbreeding as seen in a real-world wild population (Wright 1931), is used as a conservation metric and often requires a germline mutation rate to estimate. Minimum ‘healthy’ N_e_ thresholds of 50 or 500 often being used to guide wildlife management decisions (I. G. Jamieson and Allendorf 2012). Outgrowths of this have incorporated the census population size N_c_, with high N_e_/N_c_ ratios serving as an accurate indicator of high extinction risk (Wilder et al. 2023; Palstra and Fraser 2012). Knowing the germline mutation rate also has direct implications for evolutionary genomics, and can help to date the origin of a clade (Bergeron et al. 2021; Besenbacher et al. 2019) or the age of a beneficial allele (Smith et al. 2018), particularly for groups where phylogenetic estimates of *μ* are absent or problematic.

To determine if the germline mutation rate of mollusks differs from that of other animals, we measured *de novo* mutation rates in three families of the white abalone, *Haliotis sorenseni* (Fig. S1). Individuals for this study were provided by the White Abalone Captive Breeding Program, which aims to restock wild populations of this critically endangered species with aquaculture-raised individuals. *H. sorenseni* gastropod mollusk typically found in 20-60m of water along the coast of California and Mexico (T. C. Tutschulte 1976). Like many broadcast-spawning invertebrates, white abalone are highly fecund; one individual can release millions of eggs or sperm into the water column in a single spawning event, and several spawning events can occur in one year (Hobday, Tegner, and Haaker 2000). After fertilization, white abalone larvae will disperse for ∼10 days before settling on rocky substrate, after which point they will move very little, if at all, over the course of their adult lives (Leighton 1972; Lafferty et al. 2004). Growth in white abalone is slow, as sexual maturity in wild individuals occurs around four years of age with most individuals likely reproducing by six years (T. Tutschulte and Connell 1981). Individuals may live up to 30 years of age and grow to more than 20cm (Hobday, Tegner, and Haaker 2000; Andrews et al. 2013). While much is still unknown about the biology of this species, white abalone represent a unique combination of mollusk-like and vertebrate-like life history traits.

We then use our estimated GMR to resolve both recent and long-term questions in abalone conservation and evolutionary history. This species’ high fecundity in a sea of dispersive currents was once thought to buffer white abalone - and organisms with similar life histories - against overexploitation (Laura Rogers-Bennett et al. 2016) (G. Jamieson 1993). However, slow growth and overharvesting of mature adults can reduce gamete abundance and concentration to the point of total recruitment failure (Hobday, Tegner, and Haaker 2000; Stephens, Sutherland, and Freckleton 1999). Despite an estimated historical population size of 360,000 in California, white abalone populations declined precipitously during the 20th century, due to a combination of intensive fishing, disease, and this overly optimistic view of the species’ ability to recover (Laura Rogers-Bennett 2002). Given the extent of decline, white abalone was the first marine invertebrate to be listed under the U.S. Federal Endangered Species Act. A better understanding of how current population sizes stack up against historical baselines (e.g. long-term N_e_) will help guide management criteria. Additionally, the evolutionary history of white abalone and its relationship to other congeners, some of which are better studied or more robust to environmental stressors (Crosson and Friedman 2018), is largely unknown. Outlining the timescale of diversification in abalone can help set expectations for how much one species’ traits might be true of another species in this understudied group.

## Results

We observed 107 unique *de novo* mutations across the three families: nine offspring and five parents (Table 1). Only 13 (12.1%) of these mutations were inherited by two or more offspring when they had parents in common (Fig S2). After incorporating estimates of the false discovery rate (FDR) and false negative rate (FNR), 0.05 and 0.139 respectively, and the size of the callable genome (Table S1), we observe a median mutation rate of 7.99 x 10^-9^ mutations/bp/generation and a mean mutation rate of 8.60 x 10^-9^ (95% CI: 6.10 - 11.11 x 10^-9^; Fig. S3). The mean rate is faster than most arthropod per-generation rates but falls within the distribution of per-generation rates estimated for most vertebrates, plants, and the one echinoderm (Fig. 1; Wang and Obbard 2023, Popovic et al. 2024). When our rate is plotted as a function of life history traits, including approximations of generation time (∼6 years), age at sexual maturity (∼4 years), and lifespan in the wild (∼20-30 years), it remains consistent with vertebrate distributions (Fig. S4A-C; Bergeron et al. 2023, Hobday et al. 2000).

**Table 1.** Counts of observed mutations by family. **Num. mutations** = count of unique mutations across the set of offspring. Families 2 and 3, which share a mother, share two of these mutations. **Num. multi-sib mutations** = count of unique mutations found in two or more offspring of a family. **Mean rate** = Mean of the individual-level rates for each family. **Contribution** = Mutations from the paternal line (P), maternal line (M), or unknown (?) as determined by read-backed phasing only. **\*\*\***Significant parental bias; p < 0.005, chi-squared test.

| Family | Num.<br>offspring | Num.<br>mutations | Num.<br>multi-sib<br>mutations | Mean rate<br>($\times 10^{-9}$) | Contribution<br>(P / M / ? ) |
| --- | --- | --- | --- | --- | --- |
| #1 | 3 | 49 | 4 | 10.9 | 21 / 6 / 22 *** |
| #2 | 4 | 43 | 9 | 8.6 | 8 / 6 / 29 |
| #3 | 2 | 17 | 0 | 5.1 | 5 / 6 / 6 |

Using read-based haplotype phasing, we were able to attribute 29.2% of these mutations to either a mother or father. One of these families showed significant paternal bias (α), with the father contributing 21 mutations and the mother contributing only 6 (α=3.5; Table 1). The two other families, which happened to share a mother, showed roughly equal contributions from both parents (α = 1.33, 0.83). As neither shell size, weight, nor age at time of reproduction is known for all parents, it’s not possible to relate the age of each parent at spawning and to the relative contribution of mutations. We do know that two of the individuals born in captivity, the shared mother of Families 2 & 3 and the father of Family 1, were at least 19 years old at spawning. The three remaining parents were wild-caught and greater than 19 years old.

Given our estimate of a germline mutation rate for *Haliotis sorenseni*, we aimed to infer effective population size through time across *Haliotis* spp. and gather a sense of long-term baselines that could inform management of these species. To do this, we performed demographic inference and all available high coverage (>20X) whole genome sequencing data for *Haliotis* spp. Demographic analysis of five species distributed throughout the Pacific Ocean points towards large and stable effective population sizes (N_e_) over long timescales (Fig. 2A). The harmonic mean of N_e,_ summarized across roughly 1 million generations (1e4-1e6), is between 100,000 and 300,000 for all species. When we compare this value for our focal species *H. sorenseni* against the mean mutation rate, we see that values for white abalone are consistent with GMR∼N_e_ relationships in vertebrates (Fig. S4D; (Bergeron et al. 2023). *H. sorenseni* does exhibit a gradual decline over the most recent 100,000 generations, rarely surpassing an N_e_ of 50,000. The blacklip abalone - *H. rubra* - exhibits a more abrupt decline, dropping by roughly ∼90% in the past 10,000 generations. However, inferences of population size within this most recent time interval should be interpreted with caution when based on a single individual (Wilder et al. 2023). Therefore, the growth inferred for *H. cracherodii*, *H. rufescens*, and *H. laevigata* within this time interval suffers from similar limitations.

**Figure 2.**
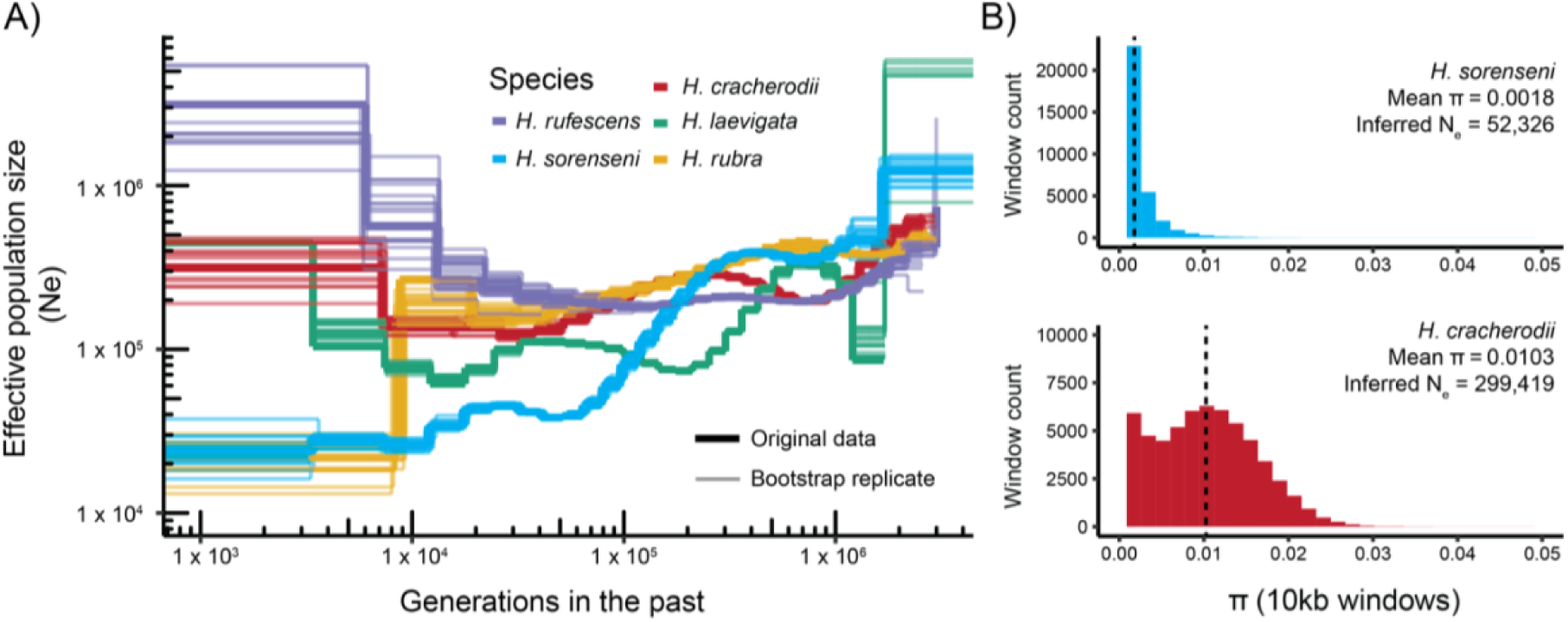
**A)** Historical effective population size (N_e_) for five abalone species as estimated by *MSMC2 ^(^Schiffels and Wang 2020).* Both original data and twenty bootstrap replicates per species are shown. **B)** Distribution of intraspecific nucleotide diversity (*π*) for species where multiple sequenced individuals were available. Effective population size is calculated from *π* = 4N_e_μ using the mean observed *π*.

These historical estimates of N_e_ based on haplotype coalescence are supported by direct calculations of N_e_ from population-level sequence diversity (*π*). We called genotypes for 11 sequenced individuals each of both *H. sorenseni* and *H. cracherodii*, the only abalone species which had sufficient population-level sequencing data. With these genotype calls, we observed intraspecific diversity (*π*) of 0.0018 and 0.0103 for *H. sorenseni* and *H. cracherodii*, respectively (Fig. 2B). By applying these values and our mean estimate of the mutation rate μ to the relationship *π* = 4N_e_μ (Nei and Tajima 1981), we obtained N_e_ values of 52,236 and 299,419. These estimates, based on population-level nucleotide diversity (Hare et al. 2011; Nadachowska-Brzyska, Konczal, and Babik 2022), closely match the historical values of N_e_ for both species (Fig. 2A).

We next applied our mutation rate to examine *Haliotis* evolution on deeper evolutionary timescales. To do this, we used genes present in the reference genomes for the above set of *Haliotis* species, the tropical abalone *Haliotis asinina* (for which a reference was also available), and the sea snail *Gibbula magus*, which served as an outgroup. We find a single highly-supported topology based on the coding sequences of 2,525 coding genes (Figure 3A). We then used a multi-species coalescent (MSC) approach to date this topology according to our germline mutation rate estimate (Tiley et al. 2020). Specifically, we used a subset of 150 clock-like genes, a mean and standard generation time of 6 and 2 years respectively (L. Rogers-Bennett, Dondanville, and Kashiwada 2004), and a mutation rate standard deviation of 3.26e-9. With this approach, we date the split between Western and Eastern Pacific abalone at 36.4 million years before present (95% CI 33.6-39.1 x 10^6^ Ma; Figure 3). Within the Western Pacific clade, the lone tropical abalone representative in this analysis, *H. asinina*, splits off from *H. rubra* and *H. laevigata* early on at 19.9 Ma. In contrast, all three Eastern Pacific abalone share a relatively recent common ancestor at 4.3 Ma (95% CI 3.9-4.7 x 10^6^ Ma), with white (*H. sorenseni*) and red (*H. rufescens*) abalone diverging only 2.2 Ma (95% CI 1.9-2.5 x 10^6^ Ma).

**Figure 3.**
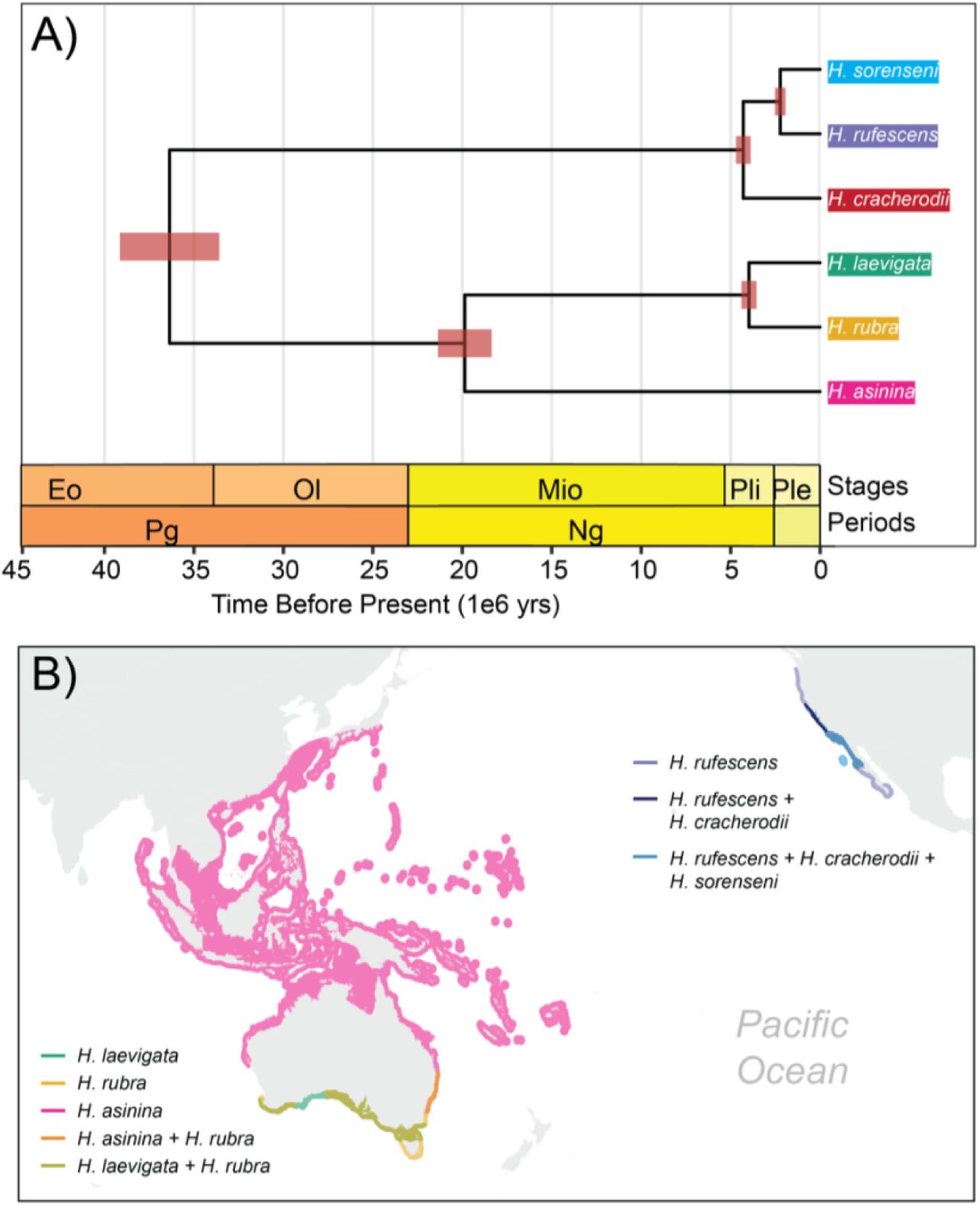
**A)** Multi-species-coalescent (MSC) phylogeny inferred with BPP (Flouri et al. 2018) and calibrated assuming μ.mean = 8.60e-9, μ.sd = 3.26e-9, g.mean = 6 and g.sd = 2, where μ is the mutation rate and g is the generation time. The tree is rooted with the outgroup *G. magus*, not shown. **B)** Distributions of the six abalone species presented in 3A. Range maps obtained from IUCN Red List.

## Discussion

We estimated the germline mutation rate of white abalone to be 8.60e-9 mutations/bp/generation. Although *H. sorenseni* is diverged from vertebrates by over 1200 Ma years of evolutionary history, the rate we estimate is similar to rates estimated previously for vertebrates with similar generation time, age at sexual maturity, and effective population size (Fig 1, Fig. S4; (Bergeron et al. 2023). Our rate is also similar to the previously estimated GMR for crown-of-thorns sea star, the only member of the phylum Echinodermata with a GMR estimate and the only other marine invertebrate and broadcast spawner for which such an analysis has been performed (9.13e-09; (Popovic et al. 2024). Our direct estimate based on pedigreed samples is more technically reliable than previous indirect estimates of mollusk mutation rates, some of which were much higher (Hoeh et al. 1996; Plough, Shin, and Hedgecock 2016) and some of which were much lower (Allio et al. 2017; A. Li et al. 2021) than ours. Altogether, these results suggest that germline mutation rates in mollusks do not deviate from the established distributions and relationships observed in other branches of the tree of life.

We observed high variance in the degree of paternal bias (α) in the contribution of *de novo* mutations to offspring. Of the three families in our analysis, two show near equal contributions from both parents (α = 1.33; 0.83), while the third family shows a clear paternal skew (α = 3.5) (Table 1). A male mutation rate bias is not unexpected in GMR studies, and the magnitude of α is known to increase with longevity (Thomas et al. 2018). The larger number of germ-cell divisions in males of many species is thought to drive a greater contribution of *de novo* mutations from fathers (Venn et al. 2014; Jónsson et al. 2017), although evidence of male bias independent of cell division number points to contributions from other processes, such as sex-specific differences in DNA damage and repair (de Manuel, Wu, and Przeworski 2022). Male and female white abalone appear to invest roughly the same amount of energy towards gonad development (T. Tutschulte and Connell 1981), but the number of sperm produced will far exceed the number of eggs produced for most abalone species (Babcock and Keesing 1999). Therefore, the greater number of germ cell divisions in male abalone could be driving the paternal bias we observe in one of our three families (α = 3.5). However, it does not explain the lack of bias in the other two families, which exhibit values of α closer to species with similar male and female reproductive input, including the crown-of-thorns sea stars (α = 0.96; (Popovic et al. 2024) and several fish species (α = 0.8; (Bergeron et al. 2023). Because the extent of male bias scales with age, it is possible that age differences between males and females in two of the three families are modifying the parental contributions. However we are unable to examine this relationship because we lack all parent ages at the time of spawning, and the wild origin of three of the five parents complicates age-size relationships measured in captivity (Mccormick et al. 2016).

Reproductive life history affects not only the degree of paternal mutation bias, but also the proportion of mutations that are shared among siblings. In our families we see that 12.1% of mutations were transmitted to multiple offspring (Table 1; Figure S2), which implies that at least 12% of the mutations we observe in the parents prior to primordial germ cell specification. Previous work has found that the sharing of germline mutations among siblings is most widespread in species with short generation times, and our results are consistent with that trend– the abalone generation time is intermediate between the generation times of mice and humans, and the proportion of mutations shared among siblings is similarly intermediate. In a previous study of mouse germline mutations, 23.9% were shared among siblings, while in humans the corresponding rate is just 4% (Lindsay et al. 2019). In the guppy *Poecilia reticulata*, which has a very short generation time of 3 months, the majority of *de novo* mutations are shared (Lin et al. 2023). Given the roughly 6 year generation time of wild white abalone, it is perhaps unsurprising that we observe a mutation rate closer to humans than that of mice or guppies.

Our inference of effective population size showed large and stable abalone populations over evolutionary timescales, including our focal species *H. sorenseni*. Over roughly 1 million generations (10^3^-10^6^), N_e_ for all five species varied between 1 x 10^5^ and 5 x 10^5^ (Fig. 2). At face value, these absolute values of N_e_ are encouraging for the future of abalone conservation. Large N_e_ populations are thought to be less susceptible to genetic drift and inbreeding depression, and N_e_ values of 50 or 500 are often cited as desirable conservation thresholds (I. G. Jamieson and ^A^llendorf 2012^)^. N_e_/N_c_, another metric of population vulnerability (Palstra and Fraser 2012; Wilder et al. 2023) further indicates population stability when considering historical abalone N_c_. For example, estimates of pre-collapse N_c_ for *H. sorenseni* and *H. cracherodii* in California are 360,000 and 3,500,000, respectively (Laura Rogers-Bennett 2002). When considering our long-term summary estimates of N_e_ (Fig. 2B), N_e_/N_c_ ratios are 0.145 for *H. sorenseni* and 0.085 for *H. cracherodii*, close to the 0.1 metric typical of healthy wild populations (Palstra and Ruzzante 2008; Frankham 1995). However, natural populations of species that have high juvenile mortality and fecundity, such as abalone, often exhibit N_e_/N_c_ ratios much lower than 0.1 (Hoban et al. 2020; Popovic et al. 2024). Uncertainties regarding life history, for example the influence of sweepstakes reproduction, make it difficult to generate clear expectations for N_e_/N_c_ in healthy populations of abalone (Hedrick 2005). The historical census size is also a source of uncertainty for abalone, but fisheries landings data (Laura Rogers-Bennett 2002), written accounts (Vileisis 2020), and population genomic data (Wooldridge et al. 2024) all depict populations of *H. sorenseni* and *H. cracherodii* as large and continuous prior to collapse.

When we compare N_e_ to contemporary population sizes, we see the degree to which recent population collapses have rendered *H. sorenseni* and *H. cracherodii* vulnerable to extinction. Both *H. sorenseni* and *H. cracherodii* are thought to exist at less than 1% of their pre-collapse abundances (Laura Rogers-Bennett 2002), meaning extreme declines in N_c_ and inflation of N_e_/N_c_. For white abalone, contemporary populations number as few as 500 and no more than 5000 (Stierhoff, Neuman, and Butler 2012; Stierhoff et al. 2014), resulting in N_e_/N_c_ as high as 100, greater than that estimated for even the most threatened mammals (Wilder et al. 2023). Black abalone show greater variation in population size. They were completely extirpated in many southern California sites (VanBlaricom et al. 2009) and are showing incipient recovery at some locations (N_c_ = 2,341 at San Nicolas Island (Kenner 2021), while at the northern end of their range declines have been less severe (Neuman, Tissot, and VanBlaricom 2010). The range in N_e_/N_c_ resulting from this variation further emphasizes the need for population-specific approaches to management. Even recovering sites like San Nicolas Island still fall in the ‘highly vulnerable’ range (N_e_/N_c_ > 100) while northern sites exhibiting minor decline are of less concern. While over-reliance of these values is not recommended, especially when decades of census data sufficiently demonstrate a species’ vulnerability, having some sense of N_e_/N_c_ - now enabled by our knowledge of the rate of input mutations - can shed light on the magnitude of vulnerability and provide important guidance for species’ recovery metrics (J. A. Robinson et al. 2022).

Our estimate of a germline mutation rate for *H. sorenseni* permitted the first time-calibrated phylogeny for the abalone genus *Haliotis* and resolved the timing of diversification of Pacific species. Despite their historical abundance, abalone are poorly represented in the fossil record (Geiger and Groves 1999). For the abalone fossils that do exist, morphological ambiguity in the fossilized shells makes it difficult to place these specimens in the context of present day diversity (Geiger and Groves 1999). Therefore, a mutation rate-based approach to phylogenetic dating is particularly suited to this system, which has yet to see an attempt at divergence dating. Our inferred topology agrees with previously reported relationships (Gruenthal and Burton 2006; Streit, Geiger, and Lieb 2006; Masonbrink et al. 2019), and we identify a common ancestor for the analyzed species at 36.4 Ma, during the late Eocene (Fig. 3). These species represent the major lineages of Pacific abalone diversity and the date is consistent with a late Cretaceous specimen found in California and an Eocene specimen from New Zealand (Estes, Lindberg, and Wray 2005; Geiger and Groves 1999). Furthermore, the estimated 4.3 Ma common ancestor of California abalone (Fig. 3) agrees with the handful of Pliocene (5.3-2.6 Ma) and much larger number of Pleistocene (2.6 Ma-11.7 ka) fossils of *H. cracherodii* and *H. rufescens* from California (Geiger and Groves 1999). Similarly, single Pleistocene fossils of *H. laevigata* and *H. rubra* from Australia are consistent with the 4.0 Ma common ancestor of these species. While the agreement between our rate-based estimates and the limited fossil record are encouraging, it should of course be recognized that these figures are subject to change as a better understanding of life history (i.e. generation time) and mutation rate variation emerges (Tiley et al. 2020). Nevertheless, these results demonstrate the utility of such an approach for closely related clades with similar constraints on fossil information.

Deriving an estimate of a species’ germline mutation rate drives at fundamental questions in biology but also contributes an essential resource for conservation and evolutionary genomics research. Here we have added to the growing understanding of how GMRs vary, finding that a significant branch of Earth’s biodiversity previously absent from the literature - in this case mollusks - exhibits mutation rate characteristics that match both empirical distributions and theoretical predictions.

## Acknowledgements

We thank the White Abalone Captive Breeding Program at the Bodega Marine Lab of UC Davis for their work in conserving this species. Brock Wooldridge was supported by the National Science Foundation Ocean Science Department (NSF–OCE) (No. 2307479).

## Author Contributions

TBW conceptualized the study with assistance from KH. TBW performed all bioinformatics analyses. SF and HC performed quality control of DNA extracts and generated all sequencing libraries. JH tracked down white abalone families and provided DNA extracts. BS provided funding for sequencing and analysis. TBW wrote the manuscript with input and approval from all authors.

## Methods

### Sample collection and DNA extraction

Samples for this study derive from the White Abalone Captive Breeding Program at the Bodega Marine Lab of UC Davis, offspring were sampled at the NOAA Southwest Fisheries Science Center in La Jolla, CA. White abalone at these facilities are bred and reared in captivity under the National Marine Fisheries Service (NMFS) Endangered Species Act (ESA) Section 10(a)(1)(A) Research Permit 14344-3R.

DNA extractions were prepared from epipodial tissue sampled from each individual. Samples from parents were obtained by excising a single epipodial tentacle from a live animal, while samples from offspring required both tentacle and epipodial fringe due to their small size and all samples came from fresh mortalities. Extractions were performed following the protocol of Gemmel and Akiyama (1996).

### Sequencing library preparation

DNA extract concentration was quantified using the Qubit dsDNA HS Assay Kit (Invitrogen) and fragment length was found with the Fragment Analyzer Genomic 50 kb DNA Kit (Agilent). Sequencing libraries were prepared following the NEBNext Ultra II FS DNA Library Prep Kit for Illumina (NEB) standard recommendations, using Y-Adapters rather than the NEBNext Adapters. All samples were diluted with 1X TE (10 mM Tris pH 8.0, 1 mM EDTA) to reach ≤100 ng inputs and incubated for 6 minutes during the enzymatic fragmentation step. Libraries were amplified for 7-8 cycles using dual unique indexes, and eluted in a final volume of 21 μL of 0.1 × TE. DNA concentration was quantified using the Qubit dsDNA HS Assay Kit (Invitrogen) and fragment length was determined using the Fragment Analyzer High Sensitivity NGS Kit (Agilent). Each library was then screened via low-coverage sequencing on an Illumina Nextseq 200 (2 x 150 bp). Libraries were then sequenced on an Illumina NovaSeq X (2 x 150 bp) with a target depth of 50x genome-wide coverage at Duke University School of Medicine’s Sequencing and Genomics Technologies Core Facility.

### Alignment and variant calling

With our raw sequencing reads, we performed initial quality control and trimmed Illumina adapters by running *fastp* (v0.23.4) with default parameters on each lane *x* sample combination of paired-end reads (Chen et al. 2018). We then merged the post-*fastp* reads for each sample. We aligned these reads to the white abalone reference genome (https://abalone.dbgenome.org/) via *bwa-mem* with *–p* to indicate interleaved paired-end fastq input, *–M* to mark short split hits as secondary for compatibility with Picard, and *-a* to output alignments of unpaired reads (H. Li and Durbin 2009). Following mapping, we marked duplicate reads in two steps using *Sentieon*: 1) *driver --algo LocusCollector --fun score_info*, then 2) *driver --algo Dedup* providing the output of step 1) with *--score_info (Kendig et al. 2019)*.

To begin variant calling, we used *Sentieon driver –algo Haplotyper --emit_mode gvcf* to create gvcf files for each individual sample. We then performed joint genotyping on this set of gvcfs using *Sentieon driver –algo GVCFtyper*, which produced cohort level variant sites across the white abalone genome. Finally, we filtered these variant sites using *GATK VariantFiltration*. We performed initial filtering on SNPs and INDELs independently, excluding SNPs with QUAL < 30.0, QD < 2.0, FS > 60.0, MQ < 40.0, MQRankSum < -12.5, ReadPosRankSum < -8.0 or SOR > 3.0 and excluding INDELs with QUAL < 30.0, QD < 2.0, FS > 200.0, ReadPosRankSum < - 20.0, SOR > 10.0. These filtering parameters were based on a combination of GATK recommendations for datasets without truth/training sets, and visual inspection of the distributions for each metric.

To validate *Sentieon* variant calls, we also called and filtered variants in parallel with *bcftools v1.13 (Danecek et al. 2021)*. First, we generated ‘pileup’ files for the set of all samples by running *bcftools mpileup --annotate FORMAT/AD, FORMAT/ADF, FORMAT/ADR, FORMAT/DP, FORMAT/SP, INFO/AD, INFO/ADF, INFO/ADR --min-MQ 20 --min-BQ 20 --max-depth 500* on all input bam files. We then piped this output into *bcftools call -m --ploidy 2* to generate vcfs for the whole set of samples. Filtering was performed using the same criteria as stated above for the GATK variants, with the exception of all FisherStrand (‘FS’) filters, which were not available via the *bcftools* method.

### Kinship matrix

To validate our sample pedigrees, we estimated a kinship matrix with *plink2 v2.00a4.4LM --make-king square –allow-extra-chr* using the set of all genome-wide biallelic SNPs (Chang et al. 2015). We visualized the resulting kinship matrix in *R 4.3.3* and confirmed that families showed the expected degree of relatedness (Fig. S1).

### Masking and determining the callable genome

Accurate estimation of germline mutation rates requires dividing the number of observed mutations by the proportion of the genome where such mutations could potentially be observed given a) genome quality and b) sequencing effort. We combined several masking approaches to determine this denominator.

We quantified mappability of the white abalone reference genome with *genmap v1.3.0 (Pockrandt et al. 2020)*. First, we indexed the genome with *genmap index*, then we determined the mappability of 150 length kmers with up to 2 mismatches using *genmap map -K 150 -E 2*. We then retained all regions where the 150x2 mappability score was less than 1.0 to create a ‘negative mappability mask’, or a list of regions with poor mappability to exclude from downstream analyses.

In addition to the above mappability mask, we also generated masks based on sequencing depth for each individual. First, we generated base-pair level resolution depth files with *samtools depth -a* for each sample, and determined the mean, median, and standard deviation of sequencing depth based on the first (largest) chromosome in the white abalone reference genome. Given these parameters, we then identified regions for each individual where read depth was either less than 20 or greater than the mean read depth plus two standard deviations. Such regions, either too low to reliably call heterozygotes or outside the standard coverage distribution for each individual, were designated as regions to mask (‘negative per-sample mask’) from subsequent analyses.

Finally, for each family, we combined the above negative mappability mask and negative per-sample masks to create a conservative set of regions to exclude from mutation rate estimation. The genome remaining after this exclusion is referred to as the ‘callable genome’, and averaged around 75-80% (Table S1).

**Table S1.** Sequencing coverage and proportion of the callable genome for each family.

| Family | Avg. seq. coverage | Pct. callable genome |
| --- | --- | --- |
| 1 | 78.48 | 76.8 |
| 2 | 62.60 | 77.4 |
| 3 | 91.95 | 79.1 |

### Mutation rate estimation

Provided variant calls from *GATK* and *bcftools* as well as callable regions of the genome, we then proceeded with mutation rate estimation. First, for each potential trio (two parents + 1 offspring), we selected variants within the callable regions that were heterozygous in the offspring and homozygous in the parents. We referred to these as candidate *de novo* mutations, and further filtered this set following the stringent criteria outlined in (Bergeron et al. 2022). Specifically, we retained mutations with a) genotype quality (GQ) greater than 60 in each member of the trio, b) no reads containing the mutation present in either parent, c) allelic balance > 0.30, and d) no occurrence in other offspring except those sharing one of the parents.

We applied this same filtering pipeline to both GATK and bcftools variant calls, and intersected those that appeared via both methods under the assumption that those appearing in only one approach were more likely to be spurious (Sendell-Price et al. 2023; Bergeron et al. 2022). Of the 2,817 de novo mutations detected via GATK and 1,023 detected via bcftools, 107 were shared.

We adjusted these raw rates for false positives and false negatives. To estimate the false discovery rate (FDR), we manually inspected the read alignments of each trio for 20 de novo mutations in IGV (P. Robinson and Others 2017). Only one of the 20 mutations did not display convincing alignment-based evidence with low alternate allelic depth and map quality < 40. Despite this result, the corresponding variant call passed our strict filters because of read realignment during *GATK*/*bcftools* variant calling. Nevertheless, we set the FDR to 5%.

To calculate the false negative rate (FNR), we first set out to identify high quality variants that would have to be heterozygous in offspring (0/1) based on the parent genotypes (e.g. 0/0 and 1/1). We then calculated what proportion of such variants, assumed to be true heterozygotes, would not pass the allelic balance, genotype quality, and depth filters listed above. This ratio was calculated on a per-child basis, and across all children exhibited a median of 0.139 and mean of 0.181. Given that this approach may overestimate the FNR because of overconfidence in parent genotypes, we opted to implement the median value of 0.139 in our FNR correction.

Finally, our reported mutation rates were calculated as:

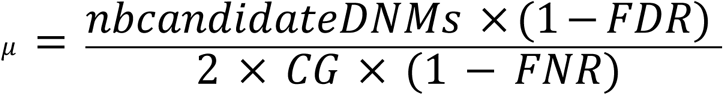

where *nbcandidateDNMs* was the count of observed de-novo mutations, and *CG* was the size, in base pairs, of the callable genome. We determined the confidence interval for the mutation rate based on the confidence interval reported by the *R* function *t.test* on the reported values of all nine offspring with estimates.

### Parent-of-origin tracing

We used read phasing in order to determine which parents contributed *de novo* mutations. To do this, we applied *POOHA* (https://github.com/besenbacher/POOHA/) to each mother-father-offspring trio of bam alignments with the options *--min-parents-GQ 60 --min-child-GQ 60 --max- marker-distance 10000 --output_variants germline*. Due to insufficient haplotype information, we were only able to trace 33 of the 107 total mutations to a parent.

### Mutation rate comparisons

In order to compare our estimated mutation rate to other published rates in multicellular eukaryotes (Fig. 1), we retrieved the set of estimates compiled by (Wang and Obbard 2023). We also obtained the corresponding phylogeny for this list of species using *TimeTree* (Kumar et al. 2017), omitting *Amphilophus* and *Marasmius oreades* when they were not located in the database. All visualizations were executed in *R* with the packages *ggtree (Yu et al. 2017)* and *aplot (Yu 2023)*.

### Analysis of polymorphisms in wild-caught *H. sorenseni* and *H. cracherodii*

To examine sequence diversity in wild populations, we first downloaded whole genome shotgun data from NCBI’s SRA database for *H. sorenseni* (n=11) and *H. cracherodii* (n=11), the only species which had multiple wild caught individuals with such data (Extended Data Table 1). We then generated variant calls for these data following the pipeline detailed in ‘Alignment and variant calling’. For both species we used their respective reference genomes (Extended Data Table 1). At the ‘GVCFtyper’ stage, in which cohort level VCFs are generated, we specified ‘-- emit_mode ALL’ in order to produce VCFs containing both invariant and variant sites. We filtered variant and invariant sites separately. For variant sites, we used the exact criteria specified for GATK filtering in ‘Alignment and variant calling’. For invariant sites, we filtered based on site quality (‘QUAL>30’) and the fraction of missing genotypes at a site (‘F_MISSING<0.25’).

We then proceeded to analyze the variant + invariant site VCFs with *pixy (Korunes and Samuk 2021)*, which estimates sequence diversity while accounting for the pitfalls in generating such estimates from heterogeneous data with high rates of missingness. We ran *pixy --stats pi -- window_size 10000*, then examined the distribution of site missingness in 10kb windows to determine filtering heuristics for downstream analysis. We reported values of *π* after retaining windows with more than 8,000 sites for *H. sorenseni* and more than 6,000 sites for *H. cracherodii*.

### Demographic inference

We reconstructed demographic histories of multiple *Haliotis* species with *MSMC2 (Schiffels and Wang 2020)* using our new mutation rate. First, we downloaded whole genome shotgun data from NCBI’s SRA database for these additional *Haliotis* species, as well as their respective reference genomes (Extended Data Table 1). We then analyzed these data following the exact pipeline detailed above in ‘Alignment and variant calling.’ Following the production of filtered variant calls, we generated two data masks: 1) reference genome mappability masks following the pipeline detailed above in ‘Masking and determining the callable genome’, and 2) sequencing depth masks following *msmc-tools* recommendations. For the latter, we used the *msmc-tools* script *bamCaller.py* on the output of *samtools mpileup -q 20 -Q 20 -C 50 -u | bcftools call -c -V indels --ploidy 2*.

With our input variant calls, reference genome mask, and sample sequencing depth mask, we then generated the *MSMC2* input with the *msmc-tools* script *generate_multihetsep.py*. We created bootstrap replicates of this input data using the *multihetsep_bootstrap.py*, specifying *-n 20 -s 5000000 --chunks_per_chromosome 10 --nr_chromosomes 20* for all species except *H. laevigata*, which had a highly fragmented genome assembly. For *H. laevigata*, we specified *-n 20 -s 100000 --chunks_per_chromosome 10 --nr_chromosomes 1000* to generate bootstrap replicates with similar characteristics as the input data. Finally, we ran *MSMC2* on the original data and all bootstrap replicates with the time segment pattern specified as *-p 25*1+1*2+1*3*. All demographic histories were then visualized in *R* and scaled by the newly estimated mutation rate of 8.60e-09 and generation time of 1.

### Phylogenomic inference

To supplement the *H. sorenseni* genome for phylogenomic analysis, we downloaded reference genome assemblies for five *Haliotis* species as well as the outgroup *G. magus* from NCBI’s RefSeq Database. (Extended Data Table 1). We subsequently annotated these genomes for BUSCO gene content using *compleasm* (v.02.6) with the *mollusca_odb10* database (Huang and Li 2023). Following this, we identified all genes that were present only as complete single copies in each of the seven taxa (six *Haliotis* + *G. magus)*, and extracted the corresponding spliced CDS nucleotide sequences from each taxon using *gffread -x (*v.0.12.8) (Pertea and Pertea 2020). Then, for each gene, we aligned the seven sequences with *mafft* (v7.526) *(Katoh and Standley 2013)* and quality trimmed the resulting alignments with *trimal -gt 0.50 -cons 50* (v.1.4.rev15) (Capella-Gutiérrez, Silla-Martínez, and Gabaldón 2009). From this set of trimmed alignments, we selected only those greater than 900bp in length, resulting in 2,525 total genes. We concatenated these genes into a single alignment with *seqkit concat* (v.0.16.1) (Shen et al. 2016) and inferred a phylogeny with *iqtree -bb 1000 -bnni -m MFP* (v.2.3.4) *(Minh et al. 2020)*. We plotted the *G. magus-*rooted tree in *R* using the package *ggtree* (v.3.10.1) (Yu et al. 2017).

Having observed 100% bootstrap support at all nodes in this initial tree, we then proceeded with inference of species divergence times under this topology. Motivated by our estimate of the germline mutation rate and a lack of obvious fossil calibrations for *Haliotis*, we opted for a fossil-free approach under the multi-species-coalescent (MSC) following principles outlined in (Tiley et al. 2020). To do this, we first estimated the extent to which each of the 2,525 genes used for tree inference evolved in clock-like fashion along the seven lineages. Specifically, we estimated the parameter ‘rate.coefficientOfVariation’ (CoV) for each gene alignment individually in *BEAST* (v.2.6.6) (Bouckaert et al. 2019). Following each BEAST analysis, we a) filtered for genes which reached an Effective Sample Size (ESS) greater than 200 for the posterior and CoV, and b) mean CoV < 0.50 and the upper and lower limits of the 95% HPD for CoV less than 1 and 0.1, respectively. This filtering resulted in 393 clock-like genes for further analysis.

With these genes and our inferred topology, we performed Bayesian estimation of divergence times under the multi-species-coalescent with *BPP (Flouri et al. 2018)*. Specifically, we applied the A00 model to a partitioned alignment of 150 randomly selected genes from the original set of 393 genes and provided the fixed topology estimated by *IQTREE.* We also set the following parameters in the *BPP* control file:

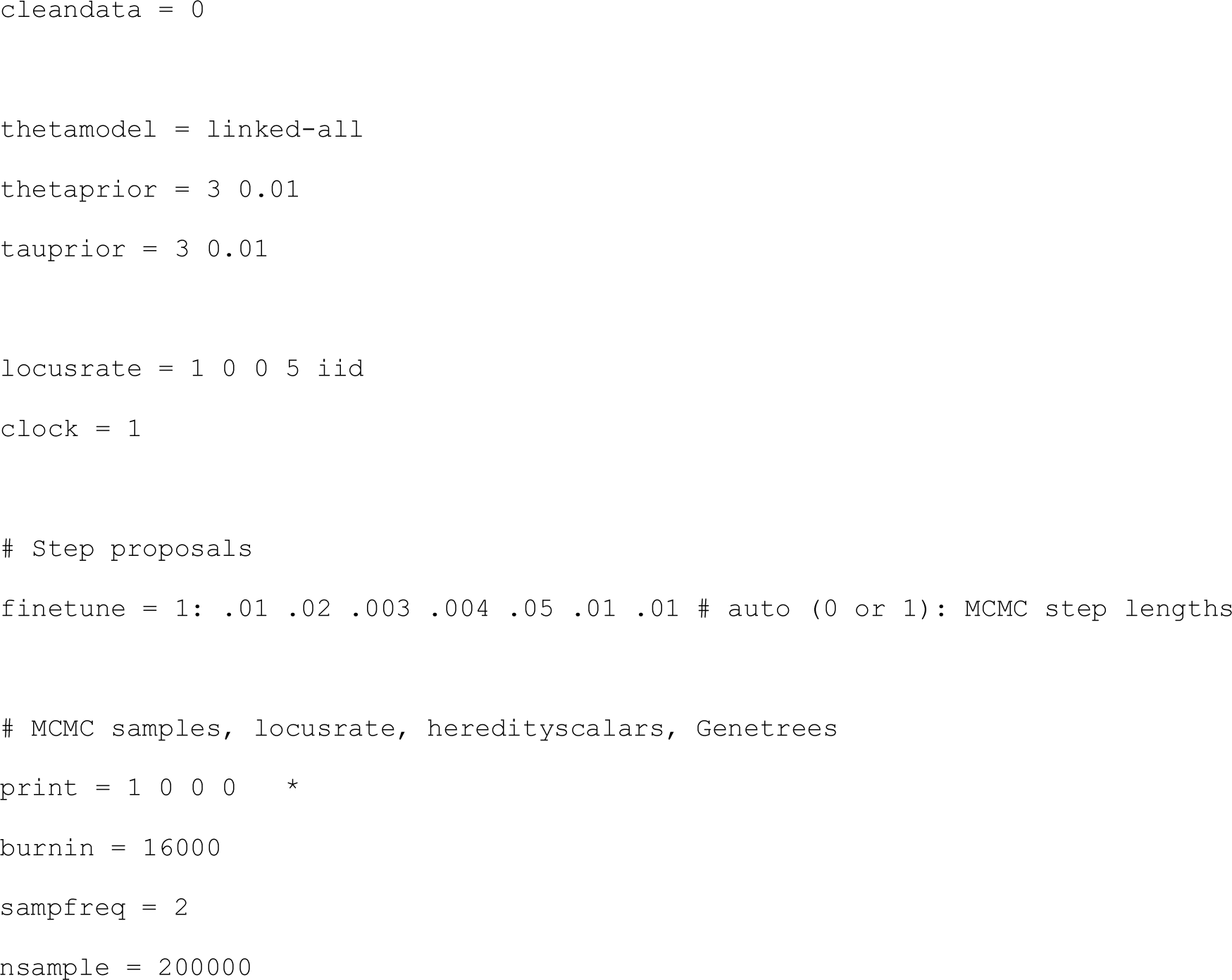

We then recalibrated branch lengths in the inferred tree to real time in R using the *msc2time.r* function from the *R* package *bppr* (). To do this, we specified the mean mutation rate *u.mean* as 8.60e-09 and the standard deviation *u.sd* as 3.26e-09 based on our findings. For lack of a precise generation time for any abalone species, we used data from growth-reproduction curves in Eastern pacific abalone like *H. sorenseni* and *H. rufescens* as a crude proxy. Specifically, we set the mean generation time *g.mean* as 6 and the standard deviation *g.sd* as 2, as most of these abalone start reproducing by at least 4 years of age and continue on into adulthood, with some evidence for reproductive senescence with age (L. Rogers-Bennett, Dondanville, and Kashiwada 2004). After using these parameters to obtain a rate for the conversion of substitution rates to real time, we rescaled all nodes, edges, and 95% confidence intervals. Finally, we reran *BPP* to confirm convergence on these parameters across independent runs.

### Species range maps

The world map was obtained from the *R* package *rnaturalearth v0.3.2 (Massicotte and South 2023)* with the function *ne_countries* and transformed to a Pacific-centered Robinson projection with the function *st_transform(st_crs("+proj=robin +lon_0=0 +x_0=0 +y_0=0 "+datum=WGS84 +units=m +pm=180 +no_defs"))* from the *R* package *sf v1.0.8 (Pebesma 2018)*. Species range shapefile were downloaded from the IUCN Red List of Species. We visualized the world map and range maps together with *ggplot2 v3.3.6*.

## Supplement

**Caption for Extended Data Table 1**. Breakdown of species and data used in this study, as well as the analysis for which they were used. **Species:** binomial nomenclature. **Sample ID:** sample shorthand label. **SRA ID:** NCBI SRA accession number, if applicable. **Reference genome:** source of genome assembly. **Mutation rate analysis**, **MSMC2**, and **PIXY:** 0 = data not used for this particular analysis, 1 = data was used for this analysis, NA = does not apply (i.e. only analyzed reference genome for said species). **Notes:** any additional sample/run information.

**Figure S1.**
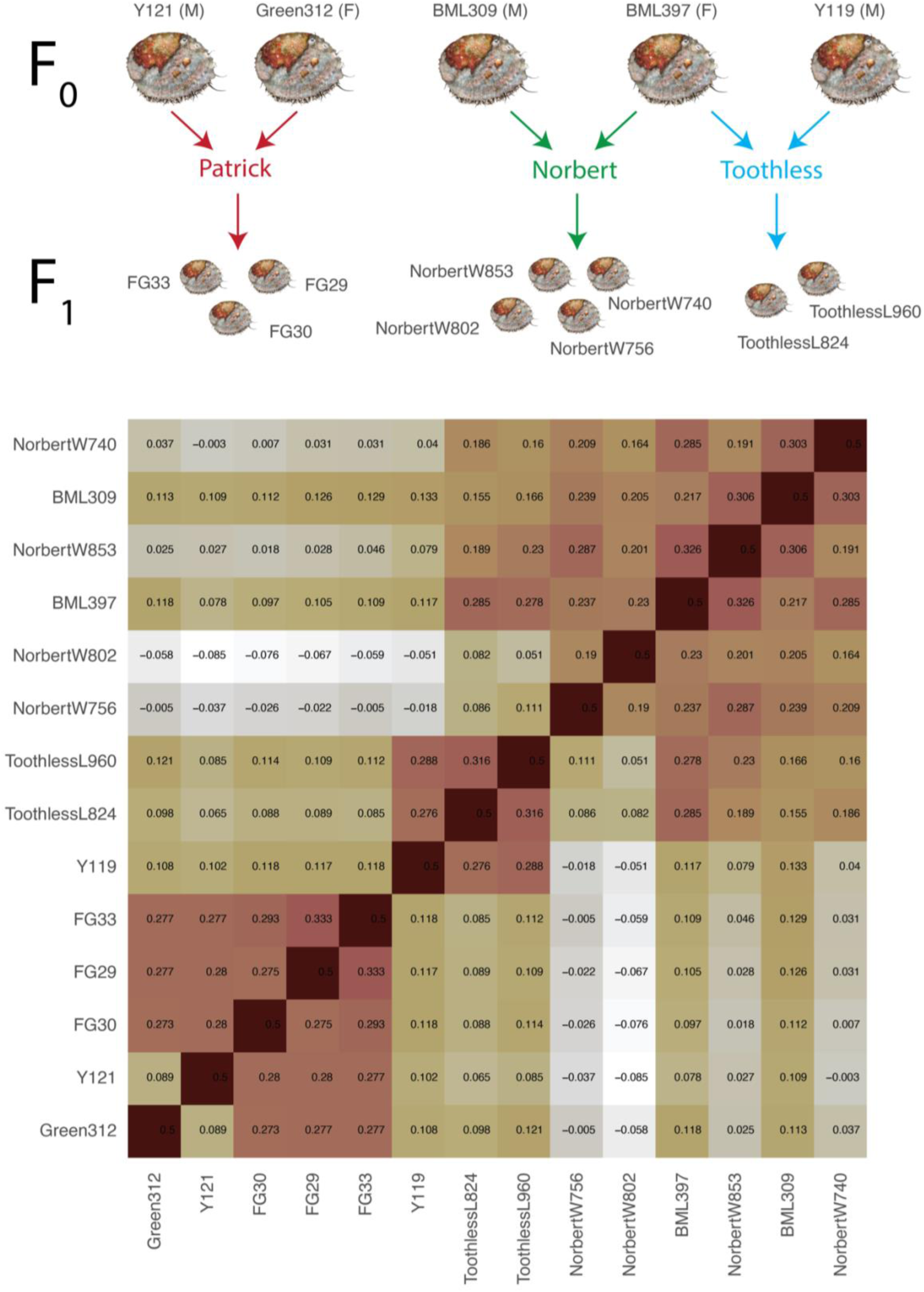
Pedigree and kinship matrix for samples included in this study. Kinship calculated with *plink2 (Chang et al. 2015)*, values are scaled so that 0.5 corresponds to perfect identity (same individual). Sexes are annotated for parents (F_0_), but unknown for offspring (F_1_).

**Figure S2.**
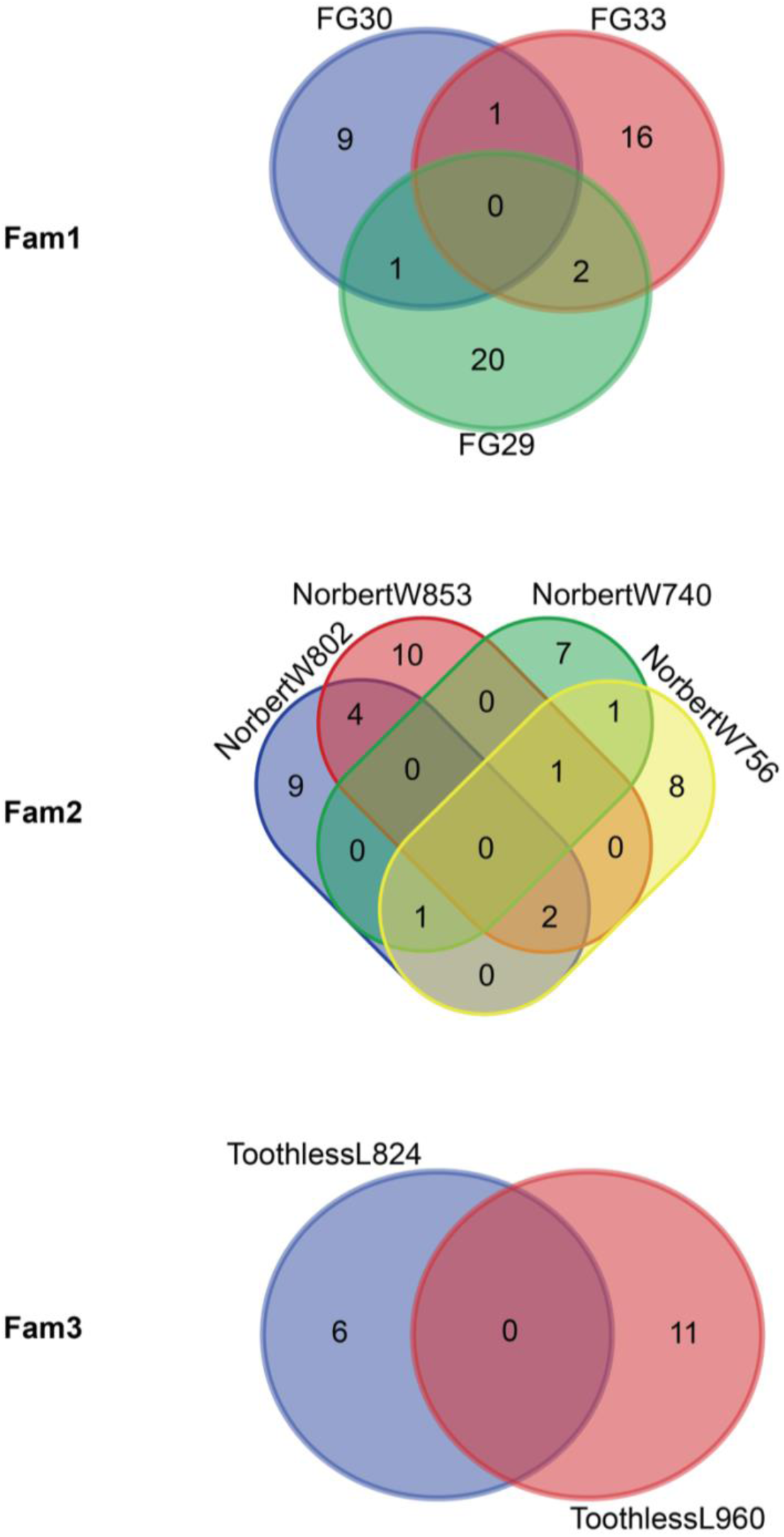
Venn diagrams showing counts of mutations unique to each individual, and shared across two or more offspring in a family. Two mutations are shared between multiple individuals of Fam2 and one member of Fam3. Thus, they are represented in the Fam2 diagram, but we do not plot all Fam2 + Fam3 intersections together for simplicity’s sake.

**Figure S3.**
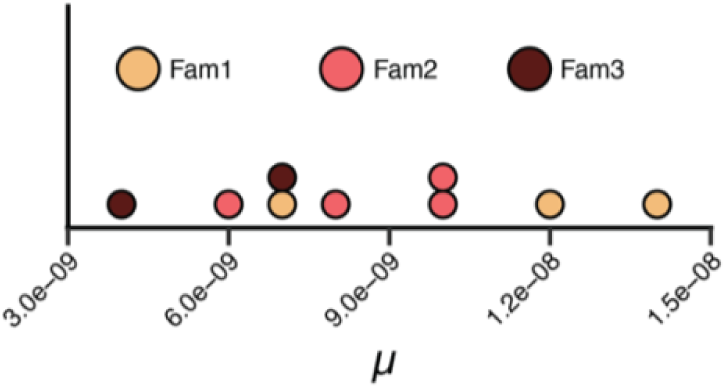
Individual-level mutation rates for all offspring included in the study.

**Figure S4.**
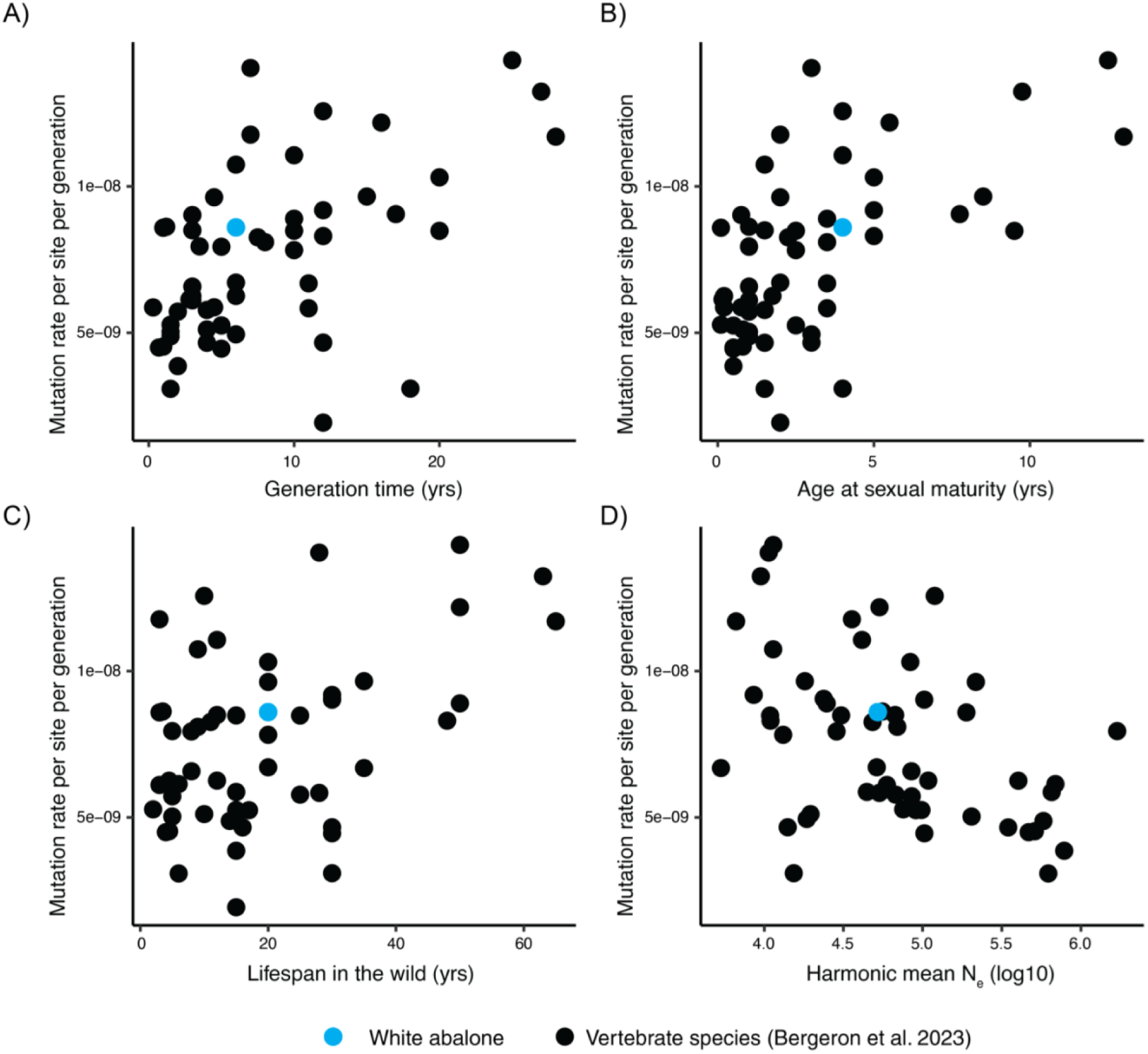
The germline mutation rate of white abalone as a function of life history traits (A-C) and demography (D). Figures reproduced from Bergeron et al. 2023. The harmonic mean N_e_ in plotted in panel D) is calculated from 5000-166,666 generations (30,000-1,000,000 abalone years assuming g = 6), following the timeframe used in Bergeron et al..

## Notes

### Competing Interest Statement

The authors have declared no competing interest.

## References

Allio, Remi, Stefano Donega, Nicolas Galtier, and Benoit Nabholz. 2017. “Large Variation in the Ratio of Mitochondrial to Nuclear Mutation Rate across Animals: Implications for Genetic Diversity and the Use of Mitochondrial DNA as a Molecular Marker.” Molecular Biology and Evolution 34 (11): 2762–72.

Andrews, Allen H., Robert T. Leaf, Laura Rogers-Bennett, Melissa Neuman, Heather Hawk, and Gregor M. Cailliet. 2013. “Bomb Radiocarbon Dating of the Endangered White Abalone (Haliotis Sorenseni): Investigations of Age, Growth and Lifespan.” Marine and Freshwater Research 64 (11): 1029–39.

Babcock, Russ, and John Keesing. 1999. “Fertilization Biology of the Abalone Haliotis Laevigata: Laboratory and Field Studies.” Canadian Journal of Fisheries and Aquatic Sciences 56 (9): 1668–78.

Bánki, O., Y. Roskov, M. Döring, G. Ower, Hernández Robles D. R., C. A. Plata Corredor, T. Stjernegaard Jeppesen, et al. 2024. “Catalogue of Life (Version 2024-07-18).” 10.48580/dgbqz.

Bergeron, Lucie A., Søren Besenbacher, Jaco Bakker, Jiao Zheng, Panyi Li, George Pacheco, Mikkel-Holger S. Sinding, et al. 2021. “The Germline Mutational Process in Rhesus Macaque and Its Implications for Phylogenetic Dating.” GigaScience 10 (5). 10.1093/gigascience/giab029.

Bergeron, Lucie A., Søren Besenbacher, Tychele Turner, Cyril J. Versoza, Richard J. Wang, Alivia Lee Price, Ellie Armstrong, et al. 2022. “The Mutationathon Highlights the Importance of Reaching Standardization in Estimates of Pedigree-Based Germline Mutation Rates.” eLife 11 (January). 10.7554/eLife.73577.

Bergeron, Lucie A., Søren Besenbacher, Jiao Zheng, Panyi Li, Mads Frost Bertelsen, Benoit Quintard, Joseph I. Hoffman, et al. 2023. “Evolution of the Germline Mutation Rate across Vertebrates.” Nature 615 (7951): 285–91.

Besenbacher, Søren, Christina Hvilsom, Tomas Marques-Bonet, Thomas Mailund, and Mikkel Heide Schierup. 2019. “Direct Estimation of Mutations in Great Apes Reconciles Phylogenetic Dating.” Nature Ecology & Evolution 3 (2): 286–92.

Bouckaert, Remco, Timothy G. Vaughan, Joëlle Barido-Sottani, Sebastián Duchêne, Mathieu Fourment, Alexandra Gavryushkina, Joseph Heled, et al. 2019. “BEAST 2.5: An Advanced Software Platform for Bayesian Evolutionary Analysis.” PLoS Computational Biology 15 (4): e1006650.

Capella-Gutiérrez, Salvador, José M. Silla-Martínez, and Toni Gabaldón. 2009. “trimAl: A Tool for Automated Alignment Trimming in Large-Scale Phylogenetic Analyses.” Bioinformatics 25 (15): 1972–73.

Chang, Christopher C., Carson C. Chow, Laurent Cam Tellier, Shashaank Vattikuti, Shaun M. Purcell, and James J. Lee. 2015. “Second-Generation PLINK: Rising to the Challenge of Larger and Richer Datasets.” GigaScience 4 (February):7.

Chen, Shifu, Yanqing Zhou, Yaru Chen, and Jia Gu. 2018. “Fastp: An Ultra-Fast All-in-One FASTQ Preprocessor.” Bioinformatics 34 (17): i884–90.

Crosson, Lisa M., and Carolyn S. Friedman. 2018. “Withering Syndrome Susceptibility of Northeastern Pacific Abalones: A Complex Relationship with Phylogeny and Thermal Experience.” Journal of Invertebrate Pathology 151 (January):91–101.

Cutter, Asher D., Richard Jovelin, and Alivia Dey. 2013. “Molecular Hyperdiversity and Evolution in Very Large Populations.” Molecular Ecology 22 (8): 2074–95.

Danecek, Petr, James K. Bonfield, Jennifer Liddle, John Marshall, Valeriu Ohan, Martin O. Pollard, Andrew Whitwham, et al. 2021. “Twelve Years of SAMtools and BCFtools.” GigaScience 10 (2). 10.1093/gigascience/giab008.

Estes, James A., David R. Lindberg, and Charlie Wray. 2005. “Evolution of Large Body Size in Abalones (Haliotis): Patterns and Implications.” Paleobiology 31 (4): 591–606.

Flouri, Tomáš, Xiyun Jiao, Bruce Rannala, and Ziheng Yang. 2018. “Species Tree Inference with BPP Using Genomic Sequences and the Multispecies Coalescent.” Molecular Biology and Evolution 35 (10): 2585–93.

Frankham, Richard. 1995. “Effective Population Size/adult Population Size Ratios in Wildlife: A Review.” Genetics Research 66 (2): 95–107.

Geiger, Daniel L., and Lindsey T. Groves. 1999. “Review of Fossil Abalone (Gastropoda: Vetigastropoda: Haliotidae) with Comparison to Recent Species.” Journal of Paleontology 73 (5): 872–85.

Gruenthal, K. M., and R. S. Burton. 2006. “Genetic Diversity and Species Identification in the Endangered White Abalone (Haliotis Sorenseni).” Conservation Genetics 6 (6): 929–39.

Hare, Matthew P., Leonard Nunney, Michael K. Schwartz, Daniel E. Ruzzante, Martha Burford, Robin S. Waples, Kristen Ruegg, and Friso Palstra. 2011. “Understanding and Estimating Effective Population Size for Practical Application in Marine Species Management.” Conservation Biology: The Journal of the Society for Conservation Biology 25 (3): 438–49.

Harrang, Estelle, Sylvie Lapègue, Benjamin Morga, and Nicolas Bierne. 2013. “A High Load of Non-Neutral Amino-Acid Polymorphisms Explains High Protein Diversity despite Moderate Effective Population Size in a Marine Bivalve with Sweepstakes Reproduction.” G3 (Bethesda, Md.) 3 (2): 333–41.

Hedgecock, Dennis, and Alexander I. Pudovkin. 2011. “Sweepstakes Reproductive Success in Highly Fecund Marine Fish and Shellfish: A Review and Commentary.” Bulletin of Marine Science 87 (4): 971–1002.

Hedrick, Philip. 2005. “Large Variance in Reproductive Success and the Ne/N Ratio.” Evolution; International Journal of Organic Evolution 59 (7): 1596–99.

Hoban, Sean, Michael Bruford, Josephine D’Urban Jackson, Margarida Lopes-Fernandes, Myriam Heuertz, Paul A. Hohenlohe, Ivan Paz-Vinas, et al. 2020. “Genetic Diversity Targets and Indicators in the CBD Post-2020 Global Biodiversity Framework Must Be Improved.” Biological Conservation 248 (108654): 108654.

Hobday, Alistair J., Mia J. Tegner, and Peter L. Haaker. 2000. “Over-Exploitation of a Broadcast Spawning Marine Invertebrate: Decline of the White Abalone.” Reviews in Fish Biology and Fisheries 10 (4): 493–514.

Hoeh, W. R., D. T. Stewart, B. W. Sutherland, and E. Zouros. 1996. “Cytochrome c Oxidase Sequence Comparisons Suggest an Unusually High Rate of Mitochondrial DNA Evolution in Mytilus (Mollusca: Bivalvia).” Molecular Biology and Evolution 13 (2): 418–21.

Huang, Neng, and Heng Li. 2023. “Compleasm: A Faster and More Accurate Reimplementation of BUSCO.” Bioinformatics 39 (10). 10.1093/bioinformatics/btad595.

Jamieson, G. 1993. “Marine Invertebrate Conservation: Evaluation of Fisheries over-Exploitation Concerns.” Integrative and Comparative Biology 33 (December):551–67.

Jamieson, Ian G., and Fred W. Allendorf. 2012. “How Does the 50/500 Rule Apply to MVPs?” Trends in Ecology & Evolution 27 (10): 578–84.

Jónsson, Hákon, Patrick Sulem, Birte Kehr, Snaedis Kristmundsdottir, Florian Zink, Eirikur Hjartarson, Marteinn T. Hardarson, et al. 2017. “Parental Influence on Human Germline de Novo Mutations in 1,548 Trios from Iceland.” Nature 549 (7673): 519–22.

Katoh, Kazutaka, and Daron M. Standley. 2013. “MAFFT Multiple Sequence Alignment Software Version 7: Improvements in Performance and Usability.” Molecular Biology and Evolution 30 (4): 772–80.

Kendig, Katherine I., Saurabh Baheti, Matthew A. Bockol, Travis M. Drucker, Steven N. Hart, Jacob R. Heldenbrand, Mikel Hernaez, et al. 2019. “Sentieon DNASeq Variant Calling Workflow Demonstrates Strong Computational Performance and Accuracy.” Frontiers in Genetics 10 (August):736.

Kenner, Michael C. 2021. “Black Abalone Surveys at Naval Base Ventura County, San Nicolas Island, California—2020, Annual Report.” Open-File Report. US Geological Survey. 10.3133/ofr20211023.

Korunes, Katharine L., and Kieran Samuk. 2021. “Pixy: Unbiased Estimation of Nucleotide Diversity and Divergence in the Presence of Missing Data.” Molecular Ecology Resources 21 (4): 1359–68.

Kumar, Sudhir, Glen Stecher, Michael Suleski, and S. Blair Hedges. 2017. “TimeTree: A Resource for Timelines, Timetrees, and Divergence Times.” Molecular Biology and Evolution 34 (7): 1812–19.

Lafferty, K. D., Behrens, G. E. Davis, P. L. Haaker, D. J. Kushner, D. V. Richards, I. K. Taniguchi, and M. J. Tegner. 2004. “Habitat of Endangered White Abalone, Haliotis Sorenseni.” Biological Conservation 116 (2): 191–94.

Launey, S., and D. Hedgecock. 2001. “High Genetic Load in the Pacific Oyster Crassostrea Gigas.” Genetics 159 (1): 255–65.

Leighton, D. L. 1972. “Laboratory Observations on the Early Growth of the Abalone, Haliotis Sorenseni, and the Effect of Temperature on Larval Development and Settling Success.” Fishery Bulletin 70 (2): 373–81.

Li, Ao, He Dai, Ximing Guo, Ziyan Zhang, Kexin Zhang, Chaogang Wang, Xinxing Wang, et al. 2021. “Genome of the Estuarine Oyster Provides Insights into Climate Impact and Adaptive Plasticity.” Communications Biology 4 (1): 1287.

Li, Heng, and Richard Durbin. 2009. “Fast and Accurate Short Read Alignment with Burrows-Wheeler Transform.” Bioinformatics 25 (14): 1754–60.

Lindsay, Sarah J., Raheleh Rahbari, Joanna Kaplanis, Thomas Keane, and Matthew E. Hurles. 2019. “Similarities and Differences in Patterns of Germline Mutation between Mice and Humans.” Nature Communications 10 (1): 4053.

Lin, Yuying, Iulia Darolti, Wouter van der Bijl, Jake Morris, and Judith E. Mank. 2023. “Extensive Variation in Germline de Novo Mutations in Poecilia Reticulata.” bioRxiv. 10.1101/2023.03.22.533860.

Lynch, Michael. 2010. “Evolution of the Mutation Rate.” Trends in Genetics: TIG 26 (8): 345–52.

Lynch, Michael, Matthew S. Ackerman, Jean-Francois Gout, Hongan Long, Way Sung, W. Kelley Thomas, and Patricia L. Foster. 2016. “Genetic Drift, Selection and the Evolution of the Mutation Rate.” Nature Reviews. Genetics 17 (11): 704–14.

Manuel, Marc de, Felix L. Wu, and Molly Przeworski. 2022. “A Paternal Bias in Germline Mutation Is Widespread in Amniotes and Can Arise Independently of Cell Division Numbers.” eLife 11 (August). 10.7554/eLife.80008.

Masonbrink, Rick E., Catherine M. Purcell, Sara E. Boles, Andrew Whitehead, John R. Hyde, Arun S. Seetharam, and Andrew J. Severin. 2019. “An Annotated Genome for Haliotis Rufescens (Red Abalone) and Resequenced Green, Pink, Pinto, Black, and White Abalone Species.” Genome Biology and Evolution 11 (2): 431–38.

Massicotte, Philippe, and Andy South. 2023. “Rnaturalearth: World Map Data from Natural Earth.” https://CRAN.R-project.org/package=rnaturalearth.

Mccormick, Thomas B., Gabriela Navas, Lorraine M. Buckley, and Christopher Biggs. 2016. “Effect of Temperature, Diet, Light, and Cultivation Density on Growth and Survival of Larval and Juvenile White AbaloneHaliotis sorenseni(Bartsch, 1940).” Journal of Shellfish Research 35 (4): 981–92.

Minh, Bui Quang, Heiko A. Schmidt, Olga Chernomor, Dominik Schrempf, Michael D. Woodhams, Arndt von Haeseler, and Robert Lanfear. 2020. “IQ-TREE 2: New Models and Efficient Methods for Phylogenetic Inference in the Genomic Era.” Molecular Biology and Evolution 37 (5): 1530–34.

Nadachowska-Brzyska, Krystyna, Mateusz Konczal, and Wieslaw Babik. 2022. “Navigating the Temporal Continuum of Effective Population Size.” Methods in Ecology and Evolution 13 (1): 22–41.

Nei, M. 1968. “The Frequency Distribution of Lethal Chromosomes in Finite Populations.” Proceedings of the National Academy of Sciences of the United States of America 60 (2): 517–24.

Nei, M., and F. Tajima. 1981. “DNA Polymorphism Detectable by Restriction Endonucleases.” Genetics 97 (1): 145–63.

Neuman, Melissa, Brian Tissot, and Glenn VanBlaricom. 2010. “Overall Status and Threats Assessment of Black Abalone (*Haliotis Cracherodii* Leach, 1814) Populations in California.” Journal of Shellfish Research 29 (3): 577–86.

Palstra, Friso P., and Dylan J. Fraser. 2012. “Effective/census Population Size Ratio Estimation: A Compendium and Appraisal.” Ecology and Evolution 2 (9): 2357–65.

Palstra, Friso P., and Daniel E. Ruzzante. 2008. “Genetic Estimates of Contemporary Effective Population Size: What Can They Tell Us about the Importance of Genetic Stochasticity for Wild Population Persistence?” Molecular Ecology 17 (15): 3428–47.

Pebesma, Edzer. 2018. “Simple Features for R: Standardized Support for Spatial Vector Data.” The R Journal. 10.32614/RJ-2018-009.

Pertea, Geo, and Mihaela Pertea. 2020. “GFF Utilities: GffRead and GffCompare.” F1000Research 9 (304): 304.

Plough, L. V., G. Shin, and D. Hedgecock. 2016. “Genetic Inviability Is a Major Driver of Type III Survivorship in Experimental Families of a Highly Fecund Marine Bivalve.” Molecular Ecology 25 (4): 895–910.

Pockrandt, Christopher, Mai Alzamel, Costas S. Iliopoulos, and Knut Reinert. 2020. “GenMap: Ultra-Fast Computation of Genome Mappability.” Bioinformatics 36 (12): 3687–92.

Ponder, Winston, and David R. Lindberg, eds. 2008. Phylogeny and Evolution of the Mollusca. Berkeley, CA: University of California Press.

Popovic, Iva, Lucie A. Bergeron, Yves-Marie Bozec, Ann-Marie Waldvogel, Samantha M. Howitt, Katarina Damjanovic, Frances Patel, et al. 2024. “High Germline Mutation Rates, but Not Extreme Population Outbreaks, Influence Genetic Diversity in a Keystone Coral Predator.” PLoS Genetics 20 (2): e1011129.

Robinson, Jacqueline A., Christopher C. Kyriazis, Sergio F. Nigenda-Morales, Annabel C. Beichman, Lorenzo Rojas-Bracho, Kelly M. Robertson, Michael C. Fontaine, et al. 2022. “The Critically Endangered Vaquita Is Not Doomed to Extinction by Inbreeding Depression.” Science 376 (6593): 635–39.

Robinson, Peter, and Others. 2017. “Integrative Genomics Viewer (IGV): Visualizing Alignments and Variants.” In Computational Exome and Genome Analysis, 233–45. Chapman and Hall/CRC.

Rogers-Bennett, Laura. 2002. “Estimating Baseline Abundances of Abalone in California for Restoration.” CalCOFI Rep. 4.

Rogers-Bennett, Laura, Kristin M. Aquilino, Cynthia A. Catton, Shelby K. Kawana, Benjamin J. Walker, Lauren W. Ashlock, Blythe C. Marshman, et al. 2016. “Implementing a Restoration Program for the Endangered White Abalone (Haliotis Sorenseni) in California.” Journal of Shellfish Research 35 (3): 611–18.

Rogers-Bennett, L., R. F. Dondanville, and J. Kashiwada. 2004. “Size Specific Fecundity of Red Abalone (Haliotis Rufescens): Evidence for Reproductive Senescence?” Journal of Shellfish Research 23 (2): 553–60.

Scally, Aylwyn, and Richard Durbin. 2012. “Revising the Human Mutation Rate: Implications for Understanding Human Evolution.” Nature Reviews. Genetics 13 (10): 745–53.

Schiffels, Stephan, and Ke Wang. 2020. “MSMC and MSMC2: The Multiple Sequentially Markovian Coalescent.” Methods in Molecular Biology 2090:147–66.

Sendell-Price, Ashley T., Frank J. Tulenko, Mats Pettersson, Du Kang, Margo Montandon, Sylke Winkler, Kathleen Kulb, et al. 2023. “Low Mutation Rate in Epaulette Sharks Is Consistent with a Slow Rate of Evolution in Sharks.” Nature Communications 14 (1): 6628.

Shen, Wei, Shuai Le, Yan Li, and Fuquan Hu. 2016. “SeqKit: A Cross-Platform and Ultrafast Toolkit for FASTA/Q File Manipulation.” PloS One 11 (10): e0163962.

Smith, Joel, Graham Coop, Matthew Stephens, and John Novembre. 2018. “Estimating Time to the Common Ancestor for a Beneficial Allele.” Molecular Biology and Evolution 35 (4): 1003–17.

Stephens, P. A., W. J. Sutherland, and R. P. Freckleton. 1999. “What Is the Allee Effect?” Oikos 87 (1): 185–90.

Stierhoff, Kevin L., Scott A. Mau, David W. Murfin, and Melissa Neumann. 2014. “White Abalone at San Clemente Island : Population Estimates and Management Recommendations.” U.S. Department of Commerce, National Oceanic and Atmospheric Administration, National Marine Fisheries Service, Southwest Fisheries Science Center. 10.7289/V5/TM-SWFSC-527.

Stierhoff, Kevin L., Melissa Neuman, and John L. Butler. 2012. “On the Road to Extinction? Population Declines of the Endangered White Abalone, Haliotis Sorenseni.” Biological Conservation 152 (August):46–52.

Streit, Klaus, Daniel L. Geiger, and Bernhard Lieb. 2006. “Molecular Phylogeny and the Geographic Origin of Haliotidae Traced by Haemocyanin Sequences.” The Journal of Molluscan Studies 72 (1): 105–10.

Sturtevant, A. H. 1937. “Essays on Evolution. I. On the Effects of Selection on Mutation Rate.” The Quarterly Review of Biology 12 (4): 464–67.

Thomas, Gregg W. C., Richard J. Wang, Arthi Puri, R. Alan Harris, Muthuswamy Raveendran, Daniel S. T. Hughes, Shwetha C. Murali, et al. 2018. “Reproductive Longevity Predicts Mutation Rates in Primates.” Current Biology: CB 28 (19): 3193–97.e5.

Tiley, George P., Jelmer W. Poelstra, Mario Dos Reis, Ziheng Yang, and Anne D. Yoder. 2020. “Molecular Clocks without Rocks: New Solutions for Old Problems.” Trends in Genetics: TIG 36 (11): 845–56.

Tutschulte, T. C. 1976. “The Comparative Ecology of Three Sympatric Abalones.” https://search.proquest.com/openview/65b982b04f70128ed41fb94229462cd8/1?pq-origsite=gscholar&cbl=18750&diss=y.

Tutschulte, T., and J. H. Connell. 1981. “Reproductive Biology of Three Species of Abalones (Haliotis) in Southern California.” The Veliger 23 (3): 195–206.

VanBlaricom, Glenn, Melissa Neuman, John L. Butler, Andrew De Vogelaere, Richard G. Gustafson, Chris Mobley, Dan Richards, Scott Rumsey, and Barbara Louise Taylor. 2009. “Status Review Report for Black Abalone.” National Marine Fisheries Service.

Venn, Oliver, Isaac Turner, Iain Mathieson, Natasja de Groot, Ronald Bontrop, and Gil McVean. 2014. “Strong Male Bias Drives Germline Mutation in Chimpanzees.” Science (New York, N.Y.) 344 (6189): 1272–75.

Vileisis, Ann. 2020. Abalone: The Remarkable History and Uncertain Future of California’s Iconic Shellfish. Oregon State University Press.

Wang, Yiguan, and Darren J. Obbard. 2023. “Experimental Estimates of Germline Mutation Rate in Eukaryotes: A Phylogenetic Meta-Analysis.” Evolution Letters 7 (4): 216–26.

Wilder, Aryn P., Megan A. Supple, Ayshwarya Subramanian, Anish Mudide, Ross Swofford, Aitor Serres-Armero, Cynthia Steiner, et al. 2023. “The Contribution of Historical Processes to Contemporary Extinction Risk in Placental Mammals.” Science 380 (6643): eabn5856.

Wooldridge, Brock, Chloé Orland, Erik Enbody, Merly Escalona, Cade Mirchandani, Russell Corbett-Detig, Joshua D. Kapp, et al. 2024. “Limited Genomic Signatures of Population Collapse in the Critically Endangered Black Abalone (Haliotis Cracherodii).” Molecular Ecology, April, e17362.

Wright, S. 1931. “Evolution in Mendelian Populations.” Genetics 16 (2): 97–159.

Yu, Guangchuang. 2023. “Aplot: Decorate a ‘Ggplot’ with Associated Information.” https://CRAN.R-project.org/package=aplot.

Yu, Guangchuang, David K. Smith, Huachen Zhu, Yi Guan, and Tommy Tsan-Yuk Lam. 2017. “Ggtree: An R Package for Visualization and Annotation of Phylogenetic Trees with Their Covariates and Other Associated Data.” Methods in Ecology and Evolution / British Ecological Society 8 (1): 28–36.

Zhang, Guofan, Xiaodong Fang, Ximing Guo, Li Li, Ruibang Luo, Fei Xu, Pengcheng Yang, et al. 2012. “The Oyster Genome Reveals Stress Adaptation and Complexity of Shell Formation.” Nature 490 (7418): 49–54.

